# Cryptic promoter activation occurs by at least two different mechanisms in the *Arabidopsis* genome

**DOI:** 10.1101/2020.11.28.399337

**Authors:** Hisayuki Kudo, Mitsuhiro Matsuo, Soichirou Satoh, Rei Hachisu, Masayuki Nakamura, Yoshiharu Y Yamamoto, Takayuki Hata, Hiroshi Kimura, Minami Matsui, Junichi Obokata

## Abstract

In gene-trap screening of plant genomes, promoterless reporter constructs are often expressed without trapping of annotated gene promoters. The molecular basis of this phenomenon, which has been interpreted as the trapping of cryptic promoters, is poorly understood. In this study, using *Arabidopsis* gene-trap lines in which a firefly luciferase (*LUC*) open reading frame (ORF) was expressed from intergenic regions, we found that cryptic promoter activation occurs by at least two different mechanisms: one is the capturing of pre-existing promoter-like chromatin marked by H3K4me3 and H2A.Z, and the other is the entirely new formation of promoter chromatin near the 5’ end of the inserted *LUC* ORF. To discriminate between these, we denoted the former mechanism as “cryptic promoter capturing”, and the latter one as “promoter *de novo* origination”. The latter finding raises a question as to how inserted *LUC* ORF sequence is involved in this phenomenon. To examine this, we performed a model experiment with chimeric *LUC* genes in transgenic plants. Using *Arabidopsis psaH1* promoter–*LUC* constructs, we found that the functional core promoter region, where transcription start sites (TSS) occur, cannot simply be determined by the upstream nor core promoter sequences; rather, its positioning proximal to the inserted *LUC* ORF sequence was more critical. This result suggests that the insertion of the *LUC* ORF sequence alters the local distribution of the TSS in the plant genome. The possible impact of the two types of cryptic promoter activation mechanisms on plant genome evolution and endosymbiotic gene transfer is discussed.

## INTRODUCITON

Gene-trap screening is a useful tool in functional genomics because it reveals the function and spatio-temporal expression profile of the genes captured by promoterless reporter constructs (Springer, 2000; Stanford *et al.*, 2001). However, gene-trap screening of plant genomes often causes unexpected expression of the constructs inserted in the intergenic genomic regions or in the reverse orientation in the coding regions, without trapping the annotated genes (Fobert *et al.*, 1994; Topping *et al.*, 1994; Ökrész *et al.*, 1998; Mollier *et al.*, 2000; Plesch *et al.*, 2000; Yamamoto *et al.*, 2003; Sivanandan *et al.*, 2005; Stangeland *et al.*, 2005). This type of enigmatic expression has been interpreted as the trapping of cryptic promoters.

Although its molecular basis is poorly understood, a cryptic promoter is presumed to be a kind of promoter whose function is not detectable unless reporter constructs are inserted just downstream of it (Fobert *et al.*, 1994; Topping *et al.*, 1994; Ökrész *et al.*, 1998; Mollier *et al.*, 2000; Plesch *et al.*, 2000; Yamamoto *et al.*, 2003; Sivanandan *et al.*, 2005; Stangeland *et al.*, 2005). Unidentified non-coding RNA (ncRNA) genes could be a source of cryptic promoters in gene-trap screening. An alternative possibility is the new occurrence of chromatin remodelling to form a chromatin structure that exhibits promoter function, just upstream of the inserted reporter genes. Although the origin and evolution of protein-coding sequences are well documented (Long *et al.*, 2003; Kaessmann, 2010; Tautz and Domazet-Lošo, 2011; Carvunis *et al.*, 2012), the mechanism via which newly emerging protein-coding sequences in the eukaryotic genome acquire transcriptional competence is less well understood.

We have been interested in the endosymbiotic evolutionary process of chloroplasts. During this process, most genes of the endosymbiotic organelle move to the host nucleus and become integrated into the nuclear gene network (Martin *et al.*, 1998; Timmis *et al.*, 2004; Matsuo *et al.*, 2005). For this functional gene transfer, the translocated genes should acquire eukaryotic promoters at their genomic integration loci. Several cases of acquisition of eukaryotic promoters by organelle-derived coding sequences by trapping pre-existing nuclear genes have been reported (Kadowaki *et al.*, 1996; Kubo *et al.*, 1999; Stegemann and Bock, 2006; Wang *et al.*, 2014). This promoter-acquisition mechanism is easy to understand, but one promoter-acquisition event will result in one disruption of pre-existing genes. Therefore an alternative mechanism might be necessary to explain how thousands of organelle-derived coding sequences have become transcriptionally active in the nucleus. To address this question, we became interested in the similarity of the promoter-acquisition event between gene-trap screening and organelle–nucleus functional gene transfer. In this respect, cryptic promoter activation is a thought-provoking phenomenon.

Recent nucleosome and transcription studies have revealed that core promoter regions in the eukaryotic genome have a specific chromatin structure: the transcriptional pre-initiation complex (PIC) and transcription start sites (TSS) occur in the nucleosome-free region (NFR), which is flanked by nucleosomes containing modified histones and histone variants (Guenther *et al.*, 2007; Li *et al.*, 2007; Cairns, 2009; Jiang and Pugh, 2009; Zhang *et al.*, 2009; Deal and Henikoff, 2011; Haberle and Stark, 2018; Andersson and Sandelin, 2020), and the downstream coding region possesses a more closed chromatin structure that represses aberrant transcription initiation within the gene body (Hennig *et al.*, 2012; Hennig and Fischer, 2013; Neri *et al.*, 2017). In this study, we scrutinized the cryptic promoters found by gene-trap screening of the *Arabidopsis* genome (Yamamoto *et al.*, 2003) and characterized them according to their chromatin-remodelling state. We found that “cryptic promoter activation” can be caused by at least two different mechanisms: one is the capturing of pre-existing promoter-like chromatin, with its inherent transcripts being hardly detectable, and the other is the entirely new formation of promoter chromatin near the 5’ end of the inserted *LUC* ORF. The latter case raises a question as to whether the inserted *LUC* ORF sequence takes part in the formation and/or localization process of the new core promoter region emerging near its 5’ end. To examine this, we performed a model promoter experiment in transgenic plants, indicating that the inserted *LUC* ORF sequence is involved in at least localization process of PIC and TSS to its 5’ proximal region. These findings for the transgenic *LUC* ORF provide new insights into a possible mechanism by which newly emerged protein-coding sequences in the plant genome acquire transcriptional competence.

## RESULTS

### Expression of the firefly luciferase (LUC) trap vector in intergenic regions

Gene-trap screening of the *A. thaliana* genome revealed that, depending on the vector design, 23% to 67% of *LUC* ORF-activation events did not depend on the capture of annotated gene promoters (Yamamoto *et al.*, 2003). To understand how a promoterless *LUC* gene (Figure 1a) can become transcriptionally active after insertion into intergenic regions, we analysed 59 *LUC*-expressing intergenic insertion lines. To simplify the analysis, eight lines were prescreened using RNA–gel blot hybridization for the discernible *LUC* transcript. Rapid amplification of cDNA 5’ ends (5’ RACE) analysis of the eight selected lines revealed that their *LUC* transcripts had the TSS in the flanking genomic regions, and not within the T-DNA inserts. From these lines, we selected YB41 and YB84 for further detailed analysis (Figure 1b) because their *LUC* transcript 5’ UTRs contained *A. thaliana* genomic sequences that were sufficiently long to examine their transcript levels, before and after the T-DNA insertion, by reverse transcription PCR (RT–PCR). The exact TSS of YB41 and YB84 were identified using the biotinylated cap-trapper method (Carninci and Hayashizaki, 1999), which showed that they were distributed 30–170 bp upstream of the T-DNA insertion sites (Figure 1c), predominantly at pyrimidine (Py) and purine (Pu) junctions (Figure S1). In WT plants, we could not detect any TSS in the corresponding genomic regions.

**Figure 1.**
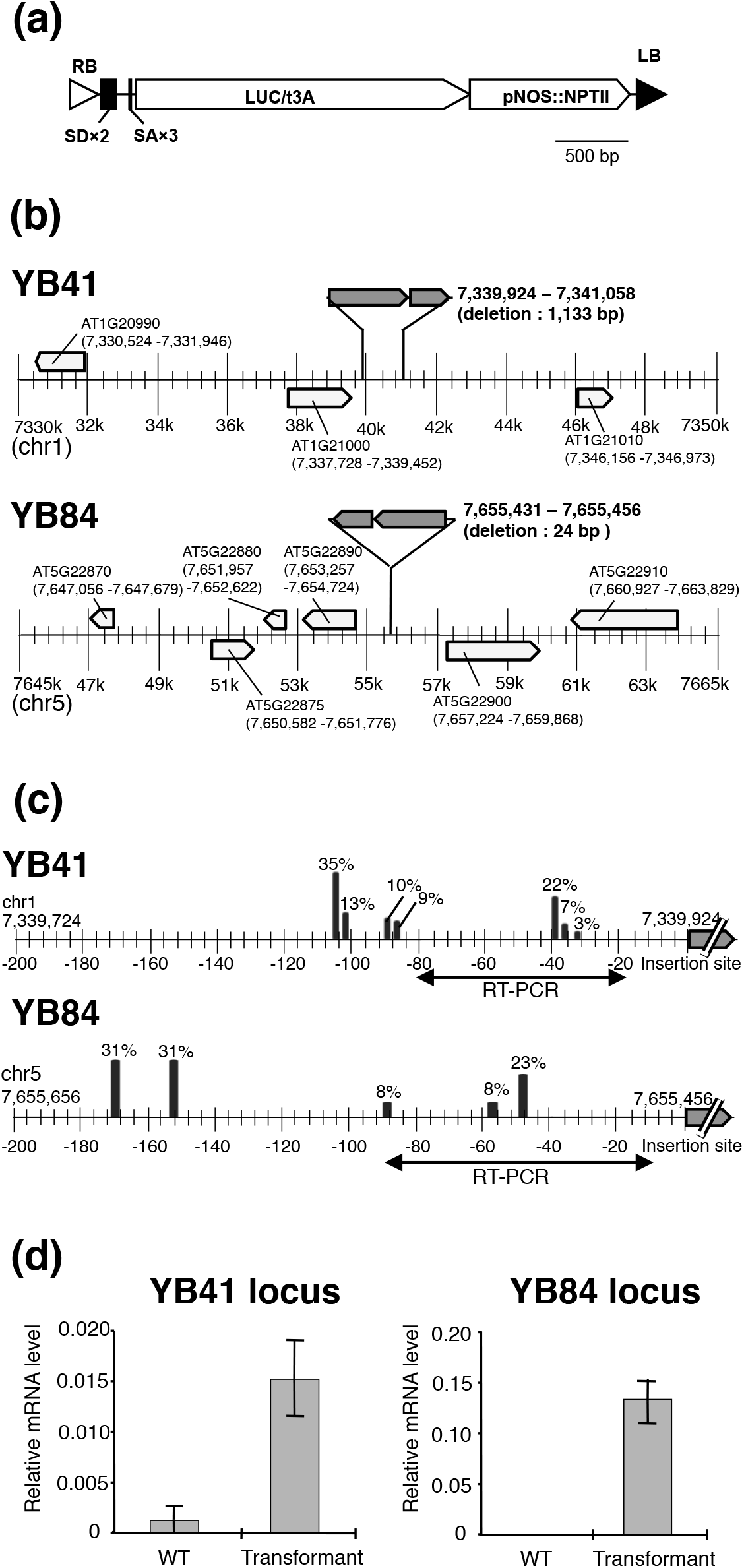
Two cryptic promoters found in the genome of *Arabidopsis thaliana* by gene trap screening. (**a**) Schematic structure of a luciferase (*LUC*)-based trap vector (Yamamoto *et al.*, 2003) containing RB (right border), LB (left border), SD (splicing donor), SA (splicing acceptor), pNOS (NOS promoter) and NPTII (kanamycin-resistance marker). (**b**) Genomic maps of the YB41 and YB84 insertion sites. (**c**) TSS distributions of the inserted *LUC* genes identified using the cap-trapper method. Genomic regions used for RT-PCR in (d) are also indicated. (**d**) Transcript levels of YB41 and YB84 integration sites determined by real-time RT-PCR analysis and normalized to the intrinsic ubiquitin (AT5G25760) mRNA level (Czechowski *et al.*, 2005). Data are means + s.d.(n=3)

We then used real-time RT–PCR to measure the transcript levels of these genomic loci before and after the T-DNA insertions normalized by the endogenous UBC (ubiquitin gene, AT5G25760) mRNA level as a reference (Czechowski *et al.*, 2005) (Figure 1d). The RNA level of the YB41 locus in transgenic plants was only 1.5% of the endogenous UBC mRNA level, with a faint signal also detected in WT plants (Figure 1d). Because no TSS was detected upstream of the YB41 locus in WT plants, this faint signal might reflect the read-through products from the upstream neighbouring gene (Figure 1b). The RNA level of the YB84 locus in transgenic plants was as high as 13% of the UBC mRNA level, whereas no RT–PCR signal was detected from WT plants (Figure 1d). These results strongly suggest that the integration of the *LUC*-trap vector at the YB41 and YB84 genomic loci (Figure 1b) caused the new occurrence of transcripts.

### Chromatin signature discrimination between “cryptic promoter capturing” and “promoter de novo origination”

We examined what occurred at the YB41 and YB84 integration sites at the chromatin level. To do this, we first prepared a custom-made DNA tiling array covering the −480 to +300 base regions of the YB41 and YB84 T-DNA insertion sites (Figure S2). We performed chromatin immunoprecipitation followed by microarray analysis (ChIP-on-chip) to examine the localization of nucleosomes containing a trimethylated form of histone H3 (H3K4me3) and a variant of histone H2A (H2A.Z), both of which are localized at core promoter regions in plants, yeast and animals (Guenther *et al.*, 2007; Li *et al.*, 2007; Cairns, 2009; Jiang and Pugh, 2009; Zhang *et al.*, 2009; Deal and Henikoff, 2011; Weber and Henikoff, 2014; Hyun et al., 2017; Haberle and Stark, 2018; Giaimo et al., 2019; Andersson and Sandelin, 2020). We also analysed the binding site of the TATA-binding protein (TBP) as a representative component of the transcriptional pre-initiation complex (PIC).

Figure 2a shows the chromatin configuration of the YB41 locus before and after the T-DNA insertion. No signals for H2A.Z, H3K4me3, TBP or TSS were detected over this genomic locus in WT plants, in which this locus is covered totally by nucleosomes containing the canonical histone H4. After T-DNA insertion, all these signals appeared and exhibited a chromatin configuration that was characteristic of the pol II TSS, as depicted in Figure 2c. The TSS occurred in the nucleosome-free region (NFR) flanked by two nucleosomes, both containing H2A.Z and H3K4me3, and TBP overlapped the 5’-flanking nucleosome (namely, –1 nucleosome relative to the NFR). The 3’-flanking nucleosome (+1 nucleosome) was located around the 5’ end of the *LUC* ORF in the T-DNA region. Because the YB41 transgenic plants used here were not homozygous, a histone H4 distribution profile similar to that of WT plants was still found in the ChIP-on-chip profile (–450 to +0). Taken together, these results indicate that a chromatin configuration that was capable of transcription initiation was newly formed at the YB41 locus after the insertion of the *LUC*-trap vector.

**Figure 2.**
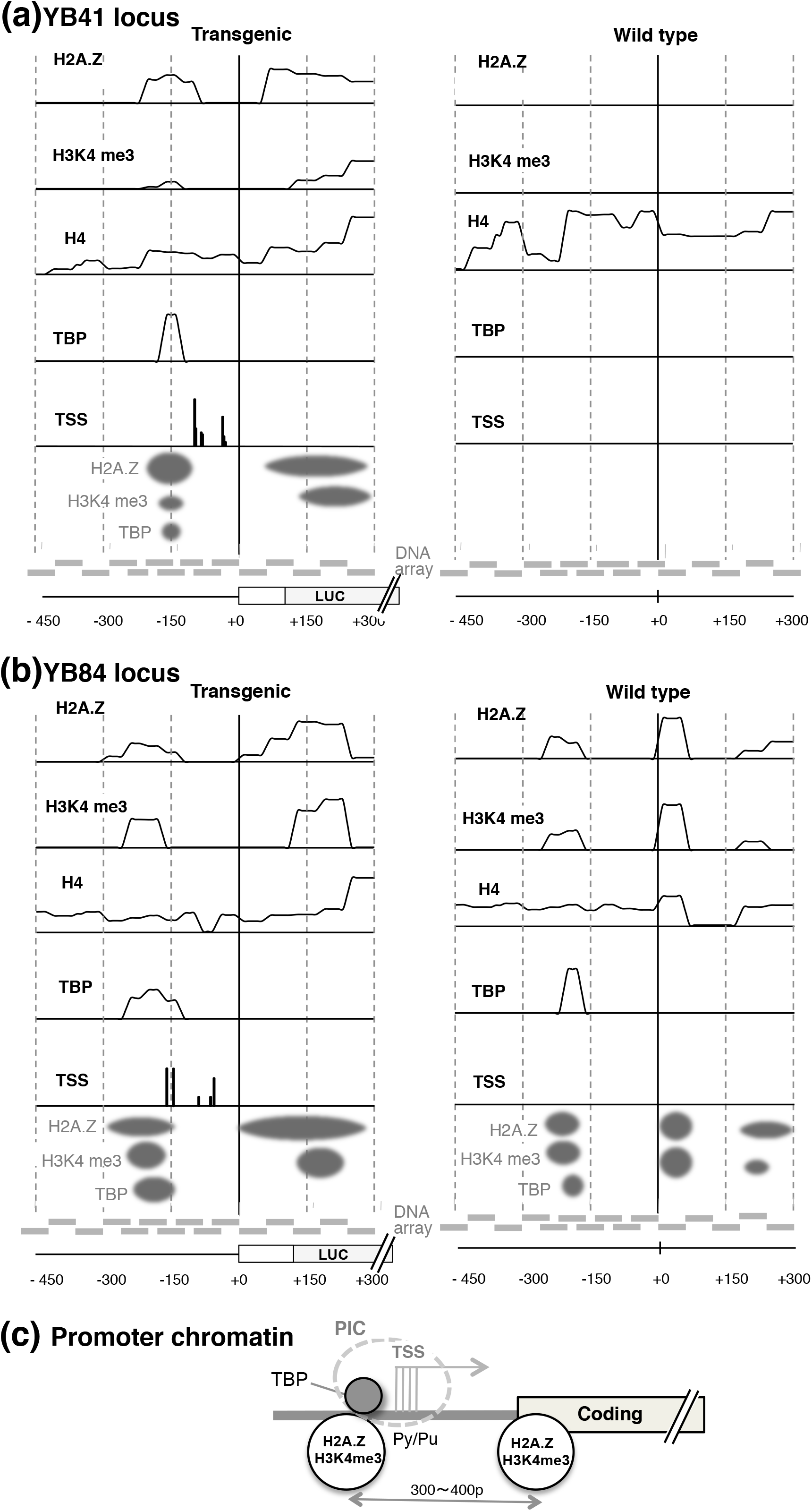
Chromatin states of the YB41 and YB84 loci before and after the insertion of a gene trap vector. (**a**) ChIP-on-chip analysis of the YB41 integration site before (right) and after (left) the T-DNA insertion. Tiling array (grey blocks) covers −480 to +300 base region relative to the genome-T-DNA junction. (**b**) ChIP-on-chip analysis of the YB84 integration site as in (a). (**c**) Schematic illustration of the chromatin structure found at the YB41 and YB84 core promoters.

The case of the YB84 locus was quite different. As shown in Figure 2b, a chromatin configuration similar to that shown in Figure 2c was already present in the WT plants. Therefore, the *LUC*-trap vector should have captured pre-existing promoter-like chromatin at this genomic locus. We did not detect any transcripts or TSS at this WT locus, which suggests that its inherent transcripts are hardly detectable because of, for example, poor stability, extremely low abundance, rapid processing and/or pausing of RNA polymerase. Even in this case, the +1 nucleosome of the transgenic plants was localized at the 5’ end of the *LUC* ORF, similarly to what was observed for YB41 (Figure 2a and 2b).

The analyses of YB41 and YB84 described above revealed that the so-called phenomenon of “cryptic promoter activation” was caused by at least two different mechanisms. To discriminate between these, we denoted phenomena such as the YB41 case as “promoter *de novo* origination” and those of YB84 as “cryptic promoter capturing”.

### The inserted LUC ORF appears to be involved in the localization of TSS to its 5’ proximal region

The finding of promoter *de novo* origination raises a question regarding the nature of its underlying mechanism. It is probable that its occurrence should depend on the individual genomic insertion sites, with as yet unidentified properties. However, it is also intriguing how the *LUC* ORF sequence is involved in this process; is it only a passively transcribed sequence, or does it have any influence on the origination and/or positioning of the TSS? To explore the latter possibility, we designed an experiment based on the following rationale (Figure 3a). In a typical protein-coding gene of plants, the core promoter region, where the PIC is formed and TSS occur, is located just upstream of the coding region and downstream of the regulatory promoter region. We attempted to separate these relationships by triplicating the core promoter segments according to three possible scenarios, as follows. (1) If the core promoter segment is an integral part of the whole promoter region, the 5’-most segment should be used preferentially (Figure 3a, i). (2) If the core promoter is an autonomous functional unit, the TSS should occur at each of the segments (Figure 3a, ii). (3) If the functional core promoter region occurs in association with the proximal coding region sequence, the 3’-most segment should be used preferentially (Figure 3a, iii).

**Figure 3.**
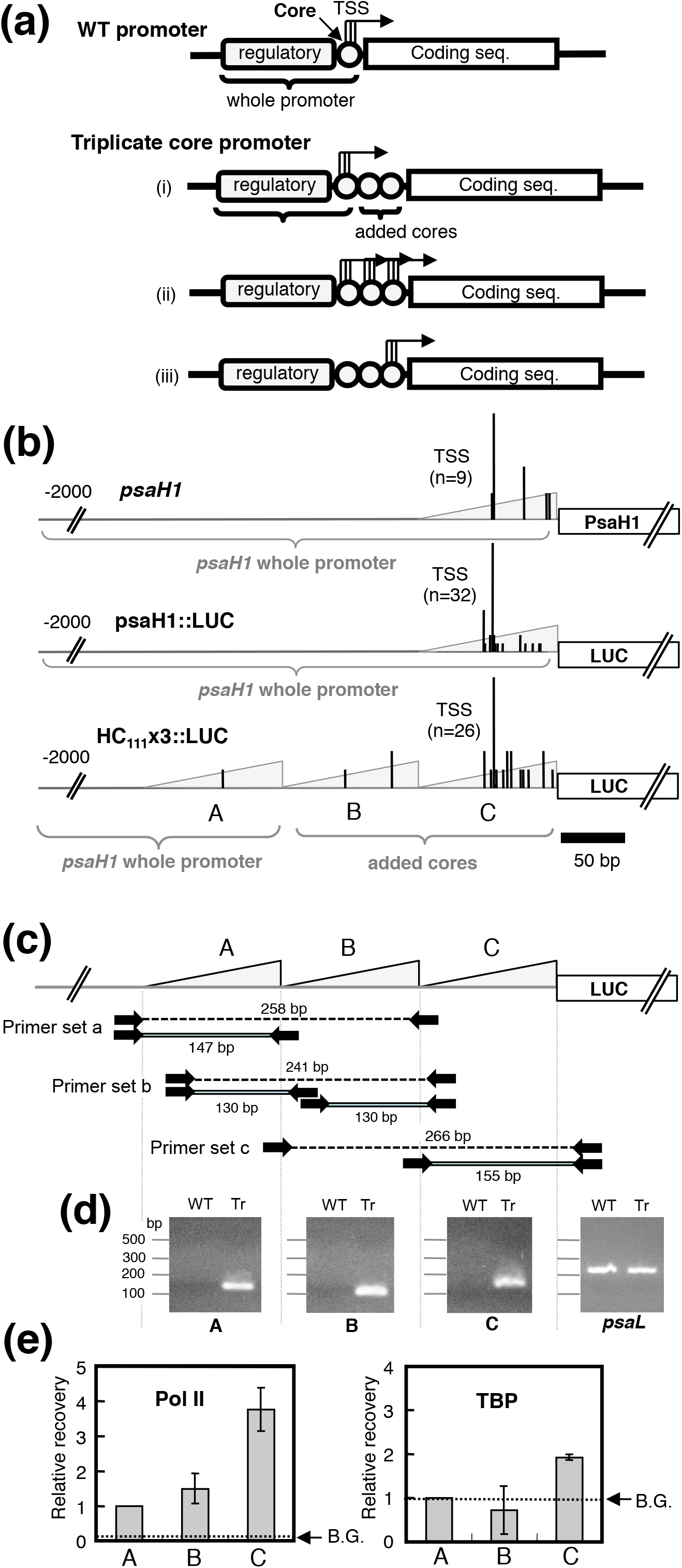
Triplicate core-promoter experiments to investigate the localization mechanism of PIC/TSS in the plant genome. (**a**) Experimental design and hypothesis. (**b**) Schematic model of the WT *psaH1* gene and chimeric promoter constructs, and TSS distributions on them. Vertical bars indicate TSS tag numbers detected at the respective genomic sites, and the heights of the peak TSSs of three genes are normalized to the same size. The total numbers of TSS tags determined for each gene are shown in parentheses as n. Triangles represent the 111 bp core promoter segment. (**c**) Strategy for the ChIP-PCR analysis of the triplicate segments. Primer sets a, b and c each can amplify both single-unit (solid line) and double-unit (dotted line) fragments of the sizes indicated. (**d**) Fine tuning of PCR conditions allowed us to amplify only single-unit fragments using primer sets a, b and c (experimental details are described in Table S2). PCR products of the chimeric core promoter segments were amplified only in the transgenic plants (Tr) and not in the wild-type plants (WT). A control primer set for an intrinsic gene (*psaL*) amplified PCR products from both WT and Tr. (**e**) ChIP-PCR analysis of the triplicate core-promoter segments A, B and C, using antibodies against pol II and TBP. ChIP signal intensities of each segment to indicate the recovery of input DNA were normalized to that of segment a, which was set at 1.0. Dotted lines indicate background levels of the control ChIP signal using mouse IgG (pol II) or rabbit IgG (TBP). Data are means ± s.d. (n=3)

To execute this experimental design with the *LUC* ORF, we triplicated the core promoter segment within the context of the promoter of the *A. thaliana psaH1* photosynthesis gene. In the endogenous *psaH1*, the TSS were distributed from –52 to –12 relative to the ATG initiation codon, with the peak TSS located at –51 (Figure 3b and Figure S3). The core promoter region of *psaH1* does not have a characteristic core promoter motif, but the TSS occur at the alteration sites of Py and Pu (Yamamoto *et al.*, 2009) (Figure S3). First, we prepared a psaH1::LUC construct in which the *psaH1* gene region (−2000 to +12) was translationally fused with the *LUC* ORF (Figure 3b, details provided in Figure S4). In the derived construct HC_111_x3::LUC, the 111 bp core promoter segment (–111 to –1) of psaH1::LUC was triplicated as direct repeats (A, B and C in Figure 3b and 3c, details given in Figure S5). The TSS selection at these artificial genes was identified using the biotinylated cap-trapper method with

RNA from transgenic plants (Figure 3b and Figures S3–S5). The TSS distribution profile of psaH1::LUC was similar to that of the WT *psaH1* gene, with a peak TSS located at –51. Surprisingly, the TSS of HC_111_x3::LUC occurred preferentially in segment C (85%), with only a small fraction observed in segments A (4%) and B (11%), even though these three segments have identical DNA sequences (Figure 3b and Figure S5). The TSS distribution profile within segment C was very similar to that observed for the core promoter region of psaH1::LUC (Figure 3b).

To confirm that the TSS occurred preferentially in segment C, we performed a ChIP–PCR analysis to examine the localization of the PIC using antibodies against pol II and TBP. Because segments A, B and C have identical DNA sequences, we discriminated between them using PCR primer sets designed at their junctions (Figure 3c). Fine-tuning of the PCR conditions allowed us to amplify a single-unit-sized DNA fragment for each primer set (Figure 3d, see methodological details in Table S2). Under this condition, the ChIP signal was quantified by real-time PCR analysis and showed that pol II and TBP were localized preferentially in segment C (Figure 3e), in accordance with the TSS distribution.

In this experiment, we did not determine the locations of nucleosomes on the chimeric promoters, because the 111 bp direct repeat sequence hinders nucleosome mapping by ChIP-on-chip or ChIP-seq analysis. However, the results obtained demonstrated that the functional core promoter region, where the PIC and TSS occur, cannot simply be determined by the sequences of its own or whole promoter region; rather, its positioning proximal to the *LUC* ORF is more critical, at least in this case.

## DISCUSSION

This study revealed that the enigmatic phenomenon of “cryptic promoter activation” was caused by at least two different mechanisms. One was the capturing of the cryptic promoter, which was the pre-existing promoter-like chromatin whose inherent transcript was hardly detectable (YB84 in Figure 1), and the other was the entirely new formation of promoter chromatin, which we denoted as promoter *de novo* origination (YB41 in Figure 1). We also analysed the third insertion line, YAB111, at the chromatin level; however, during this analysis, we became aware that this line trapped a microRNA gene, miR398 (Sunkar *et al.*, 2012). We also found that the *LUC* insert of YB84 line located in the vicinity of a transposable element (AT5TE27670). Because some transposable elements have promoter activities (Feschotte, 2008), it may be responsible for the transcriptional activation of YB84 line, although we did not observe the evidence of active transcription of this element. These imply that some portion of the cases that are currently thought of as “cryptic promoter capturing” may be reclassified as the capturing of some ncRNA genes or the other non-coding elements. In respect to this, the density of genetic elements other than protein-coding genes in the plant genome deserves further attention. Now we are proceeding with large-scale study focusing on the relative frequency of the cryptic promoter capturing and promoter *de novo* origination (Satoh and Hata *et al.*, 2020).

Our finding of “promoter *de novo* origination” in YB41 raises the question of how it occurs. Because a promoter-specific chromatin structure was not found in the YB41 locus of wild-type plants, it seems likely that the insertion of the *LUC*-trap vector sequence triggered chromatin remodeling to form the core promoter chromatin. The underlying mechanism could be investigated from two angles: the properties inherent to individual genomic insertion sites, and the possible roles of the inserted sequences. Regarding the first aspect, more examples are needed to analyse the common properties of the chromosomal integration sites.

Relevant to the second aspect, the triplicate core promoter experiment (Figure 3) revealed three intriguing properties of the transcription initiation of *LUC* chimeric genes. First, the transcription initiation region could not be determined only by the promoter sequences. Second, the sequences located downstream of the TSS, in this case the *LUC* ORF sequence, appeared to be involved in the determination of the transcription initiation region. Third, once the transcription initiation region was fixed, fine TSS distribution within the region was determined by the region’s sequence (Figure 3b and S5). Taking these hierarchical properties of the TSS determination of the *LUC* chimeric gene into consideration, it is very likely that the inserted *LUC* ORF sequence is involved in the positioning process of the newly emerged transcription initiation region proximal to its 5’ end. In this regard, it is quite intriguing that the +1 nucleosomes (3’-flanking nucleosome of the NFR) of YB41, YB84 and YAB111 are all located at the 5’ end of the *LUC* ORF, thus within the inserted T-DNA region. This suggests that the inserted *LUC* ORF sequence provides a suitable site for fixing the +1 nucleosome. The position and remodelling of the +1 nucleosome is important for the nucleosome landscape of the promoter and coding regions (Mavrich *et al.*, 2008; Jiang and Pugh, 2009; Möbius and Gerland, 2009; Valen and Sandelin, 2011; Lenhard *et al.*, 2012, Klemm *et al.*, 2019). The NFR is generally a one-nucleosome-wide region located between the –1 and +1 nucleosomes, and H3K4me3 of these nucleosomes is thought to interact with TAF of TFII-D to localize the PIC (Lenhard et al., 2012; Lauberth *et al.*, 2013). Therefore, the mechanism via which the +1 nucleosome is localized at the 5’ end of the inserted *LUC* ORF deserves further attention. In yeast, histone chaperons FACT and Spt6 contribute to the promoter-specific deposition of H2A.Z by selectively preventing the accumulation of H2A.Z within the gene bodies (Jeronimo *et al.*, 2015). Similar mechanism by which plant genome localizes H2A.Z in the 5’ proximal region of the gene bodies may operate promoter-specific chromatin remodeling (Verbsky and Richards, 2001; Choi *et al.*, 2007; Sura *et al.*, 2017; Potok *et al.*, 2019). This possibility requires further examination.

Although the molecular mechanism underlying the “cryptic promoter capturing” and “promoter *de novo* origination” remains to be analysed, the discovery of these transcriptional activation mechanisms provides important clues to elucidate how newly emerged protein-coding sequences in the plant genome acquire transcriptional competence. For example, in the promoter-acquisition process of endosymbiotic gene transfer from the organelle to the nucleus, these new activation mechanisms will leave negligible traces of the activation process on the transcriptionally activated genes, and yield little damage to the pre-existing nuclear gene network compared with the conventional model of foreign-gene trapping of pre-existing nuclear gene promoters (Kadowaki *et al.*, 1996; Kubo *et al.*, 1999; Stegemann and Bock, 2006; Wang *et al.*, 2014). We speculate that the expression level of the coding sequences that are activated by these mechanisms might generally be as low as the basal transcription level; however, from the evolutionary viewpoint and timescale, once proto-genes (Carvunis *et al*, 2012) or young genes obtain basal transcription activity, they may evolve and acquire better promoter context and elements via subsequent natural selection. We expect that this speculated mechanism will provide a new explanation for the promoter-acquisition process of the following cases: evolution of the protein-coding sequences emerging in response to, for example, stochastic changes in the genome sequences or exon shuffling (Long *et al*, 2003; Kaessmann, 2010; Tautz and Domazet-Lošo, 2011; Carvunis *et al.*, 2012,), horizontal gene transfer (Keeling and Palmer 2008; Syvanen, 2012; Soucy *et al.*, 2015; Husnik and McCutcheon, 2018) and organelle-to-nucleus DNA flux (Timmis *et al.*, 2004; Matsuo *et al*, 2005; Bock, 2017).

Based on the cryptic promoters found in gene-trap screenings of the plant genome, this study led us to find two activation mechanisms of cryptic promoters, and to a speculation regarding the potential impact of these phenomena on the plant genome evolution. Although this study was intensive regarding both time and effort, its final output included only a few gene examples. To extend this study regarding both the number of examples and the depth of the molecular analysis, we are improving its general experimental design to achieve a high-throughput analysis, which will be described elsewhere (Satoh and Hata *et al.*, 2020).

## MEXPERIMENTAL PROCEDURES

### Gene-trap plant lines

*LUC*-expressing intergenic insertion lines were screened from the gene-trap lines of *Arabidopsis thaliana* (Yamamoto *et al.*, 2003).

### Northern hybridization analysis

Northern hybridization was performed as described previously (Matsuo and Obokata, 2002). The hybridization probe was prepared from the *LUC* gene using the primer pairs LUCpr1 and LUCpr2 (Table S1).

### 5’ RACE analysis

5’ RACE was performed using the 5’-full RACE Core Set (TKR6122, TaKaRa) according to the manufacturer’s instructions. The primers that hybridized within the luciferase-coding sequence were L-RT, L-S1, L-A1, L-S2 and L-A2 (Table S1).

### Determination of the TSS

The TSS were identified in the WT (col-0) and transgenic (YB41, YB84, psaH1::LUC and HC_111_x3::LUC) seedlings of *A. thaliana* grown on MS agar plates for 10 days using the biotinylated cap-trapper method (Carninci and Hayashizaki, 1999). Total RNAs were extracted from the aerial parts of the seedlings, and cDNAs were synthesized using the following primers: for the transgenic plants, an equimolar mixture of polyT primer (5’-NT_20_-3’) and the *LUC*-gene-specific reverse primer, LUCR2 (Table S1), was used; for the WT plants, an equimolar mixture of the polyT primer (5’-NT_20_-3’) and YB41wt or YB84wt primers (Table S1), which hybridize downstream of the YB41 and YB84 junction sites, respectively, was used. The resultant cDNAs were purified using the biotinylated cap-trapper method, to give full-length cDNAs, which were subsequently ligated with a synthetic linker, GN5 (Table S1), at their 3’ ends, and amplified by nested PCR with primers corresponding to the linker and coding-region sequences. The PCR products obtained were cloned into the T-Vector (pMD20, TaKaRa), and the plasmid DNAs obtained were sequenced using an ABI PRISM 3100 Genetic Analyzer (Applied Biosystems).

### Real-time RT–PCR analysis of the YB41 and YB84 trap lines

First-strand cDNAs were synthesized from 1 μg of total RNA using ReverTra Ace (TOYOBO) and an oligo dT primer (18-mer). Real-time PCR experiments were performed with the Thunderbird® qPCR Mix (TOYOBO) and the Eco Real-Time PCR System (Illumina). The primers and thermal cycling conditions used for real-time PCR analysis are summarized in Table S2.

### ChIP-on-chip analysis

WT (col-0) and transgenic (YB41 and YB84) *A. thaliana* seedlings grown on MS agar plates under continuous white light for 10 days were subjected to cross-linking and chromatin isolation as described by Saleh *et al.* (Saleh *et al.*, 2008), with modifications. The isolated nuclei were suspended in MN digestion buffer (500 U/mL of micrococcal nuclease, 3 mM CaCl_2_, 5 mM MgCl_2_, 60 mM Kill, 15 mM NaCl, 0.25 M sucrose and 50 mM HEPES; pH 7.5) and incubated at 37 °C for 8 min, and the digestion was stopped by the addition of a 0.25 volume of nuclear lysis buffer (150 mM NaCl, 10 mM EDTA, 1% SDS, 0.1% sodium deoxycholate, 1% Triton X-100, 50 mM HEPES; pH 7.5). After the addition of 10 volumes of ChIP dilution buffer (50 mM HEPES, pH 7.5, 150 mM NaCl, 0.0875% sodium deoxycholate, 0.875% Triton X-100, 1 mg/mL pepstatin A and 1 mg/mL aprotinin), the mixture was centrifuged at 14,000 × *g* for 10 min and the supernatants were subjected to chromatin immunoprecipitation according to the method of Kimura *et al.* (Kimura *et al.*, 2008), with slight modifications. Antibodies used in this study were described below. The immunoprecipitated DNAs (IP DNAs) obtained were blunted with T4 DNA polymerase, ligated to the annealed products of linker1 and linker2 (Table S1) and PCR amplified with linker1. The amplified IP DNAs were labelled with the BioPrime Array CGH Labeling System (Invitrogen) according to the manufacturer’s instructions. One microgram each of IP DNA and control Input DNA were labelled with cy5 and cy3, respectively, mixed and precipitated with ethanol. The pellet was dissolved in 10 μL of hybridization buffer (10% formamide, 0.02 g dextran sulfate, 3× SSC, 20 μg yeast tRNA, 4% SDS, 20 μg human cot-1 DNA (Invitrogen)), dropped onto a custom-made DNA chip (NGK Insulators Ltd.) (Figure S1), covered with a coverglass and incubated at 42 °C for 24 h in a hybridization chamber. The DNA chip was designed to contain a 60-mer tiling array covering –480 to +300 relative to the genome–T-DNA (YB41 and YB84) junctions and their corresponding WT genomic regions. After incubation, the DNA chip was washed once with 2× SSC with 0.1% SDS at 30 °C for 5 min, once with 2× SSC with 50% formamide (pH 7.0) at 30 °C for 15 min, once with PN buffer (0.1 M NaH_2_PO_4_/Na_2_HPO_4_, pH 8.0, 0.1% NP-40) at 30 °C for 30 min, and finally with 2× SSC at room temperature for 5 min. The hybridization signals were analysed on a GenePix 4000B scanner using the GenePix Pro 4.0 software (both from Axon Instruments).

### Chimeric promoter constructs and plant transformation

The promoter region (–2000 to +12 relative to the ATG initiation codon) of *psaH1* (AT3G16140.1) of *A. thaliana* was translationally fused with the firefly *LUC* gene and cloned into pPZP221 to obtain the psaH1::LUC construct (sequence details are given in Figure S4). HC_111_x3::LUC was generated by triplicating the 111 bp segment (–111 to –1) of psaH1::LUC (sequence details are given in Figure S5). *Agrobacterium-mediated* transformation of *A. thaliana* col-0 was performed as described previously (Yamamoto *et al.*, 2003).

### ChIP–PCR analysis of the triplicate core promoter construct

Cross-linking treatment of *A. thaliana* seedlings was performed as described above. Chromatin was isolated according to the method of Gendrel *et al.* (Gendrel *et al.*, 2005) and was suspended in nuclear lysis buffer (50 mM Tris-HCl (pH 8.0), 10 mM EDTA, 1% SDS, 1 mM PMSF, 2 μg/mL pepstatin A and 2 μg/mL aprotinin). After the addition of nine volumes of ChIP dilution buffer (50 mM Tris-HCl (pH 8.0), 167 mM NaCl, 1.1% Triton X-100, 0.11% sodium deoxycholate, 1 mM PMSF, 1 μg/mL pepstatin A and 1 μg/mL aprotinin), chromatins were fragmented to 50–500 bp, with a peak at 200 bp, by sonication using a UD-201 ultrasonic disruptor (Tomy Seiko). Chromatin immunoprecipitation was performed essentially as described above, using antibodies against pol II and TBP. The IP DNAs obtained were subjected to real-time PCR analysis, as summarized in Table S2.

### Antibodies

The anti-*A. thaliana* TBP rabbit polyclonal antibodies were prepared using the synthetic peptides TBP1a (N-PVDLSKHPSGIVPTL-C), TBP1b (N-GFPAKFKDFKIQNIV-C) and TBP1c (N-ENIYPVLSEFRKIQQ-C). These three peptides were injected into different rabbits. The anti-*A. thaliana* H2A.Z rabbit polyclonal antibodies were prepared according to the method of Deal *et al.* (Deal *et al.*, 2007): equal amounts of synthetic peptides representing the N-termini of HTA9 (At1g52740) and HTA11 (At3g54560) were mixed and injected into rabbits. The anti-H3K4me3 mouse polyclonal antibody was described previously (Kimura *et al.*, 2008). A mouse monoclonal antibody (8WG16) against RNA polymerase II CTD repeats SPTSPS was purchased from Abcam. Normal mouse IgG (sc-2025) and rabbit IgG (sc-2027) were purchased from Santa Cruz Biotechnology, Inc.

## ACKNOWLEDGEMENTS

We thank M. Sugiura, T. Yoshitsugu, T. Wakasugi, M. Tachikawa and A. Takabayashi for discussions and encouragement; S. Shirakawa, K. Hasegawa, Y. Yasui and M, Ohtani for assistance with experiments; and S. Kushnir for critical reading of this manuscript. This work was supported in part by grants from the MEXT Japan and JSPS (grant numbers 22370002, 23125512, 23657040, 23117006 and 25650131 to JO), from Sekisui Chemical Co. Ltd., and the Strategic Research Funds at Kyoto Prefectural University.

## SUPPORTING INFORMATION

Additional supporting information is found in the online version of this article.

**Figure S1.**
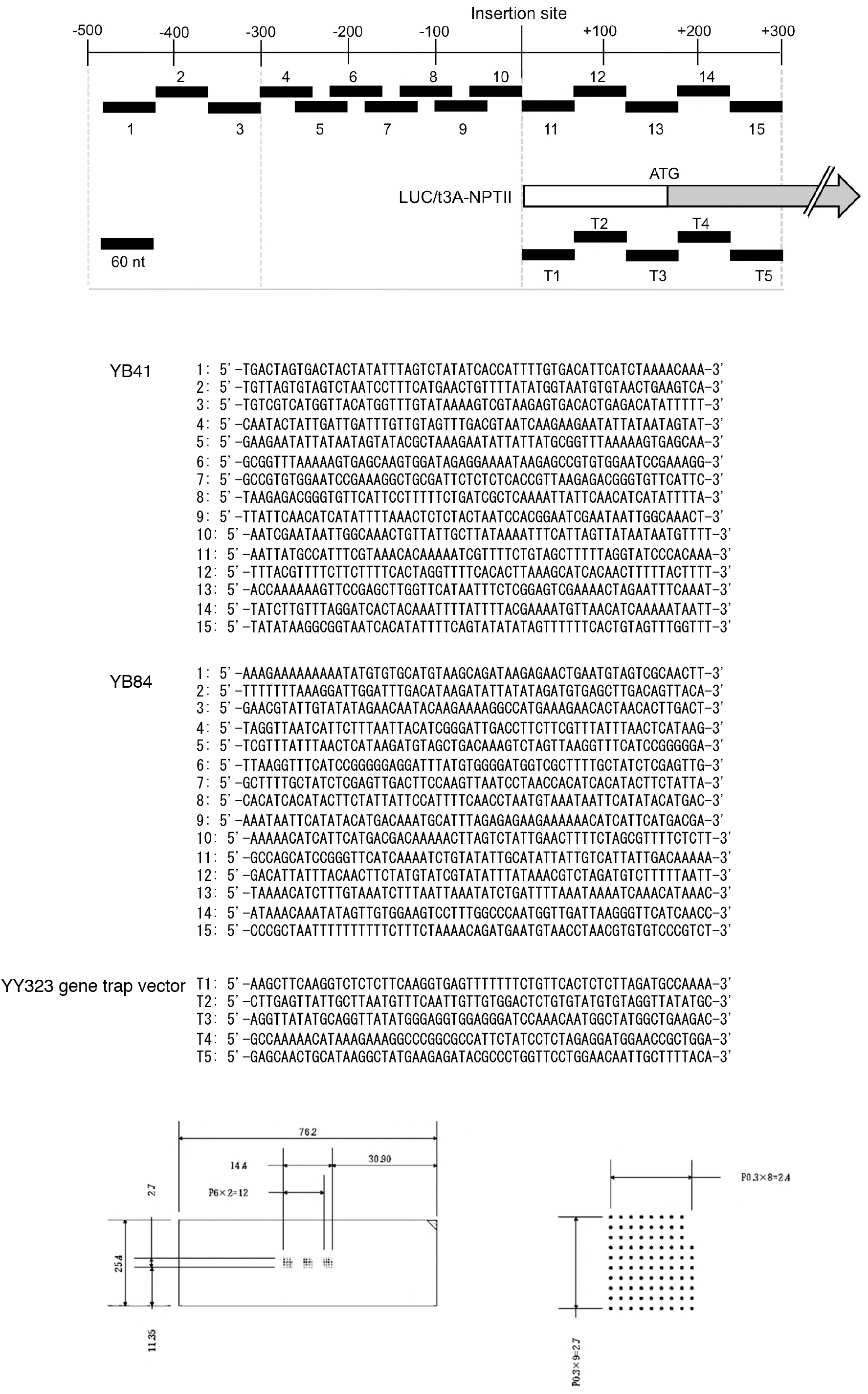
Custom-made DNA tiling array. The top panel shows the layout of DNA fragments 1—15, and T1—T5, on the schematic gene map. Nucleotide sequences of the synthetic 60-mer DNA fragments are shown in the middle panel. The bottom panel shows the spotting layout on the slide glass.

**Figure S2.**
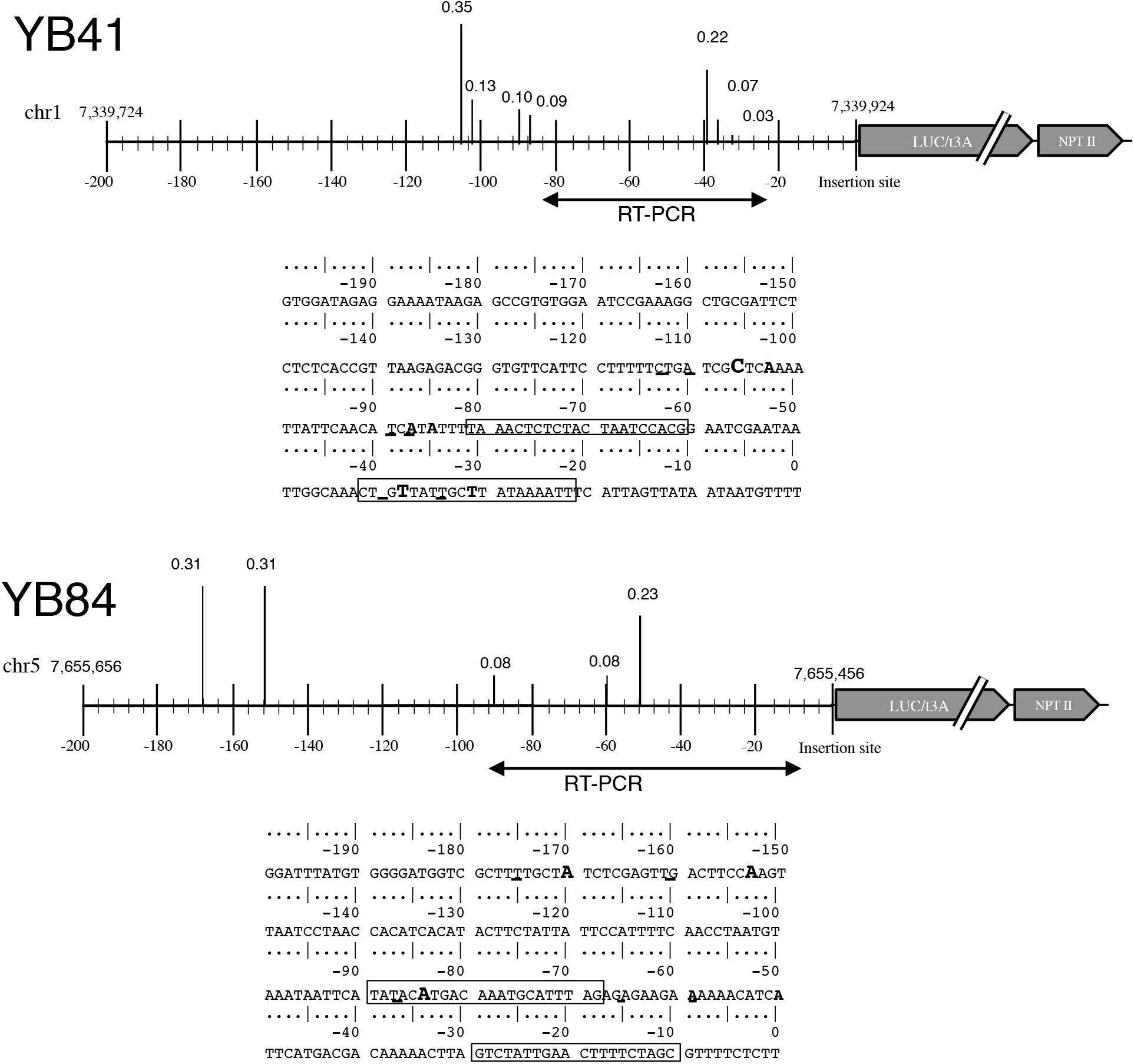
Detailed sequence and TSS distributions of the YB41 and YB84 loci. Vertical bars on the sequence map show the distribution profile of TSSs, with the total sum of TSSs as 1.0. Bold letters in the nucleotide sequences indicate TSSs. Boxes indicate the sites of PCR primers used for RT-PCR.intrinsic *psaH1* gene

**Figure S3.**
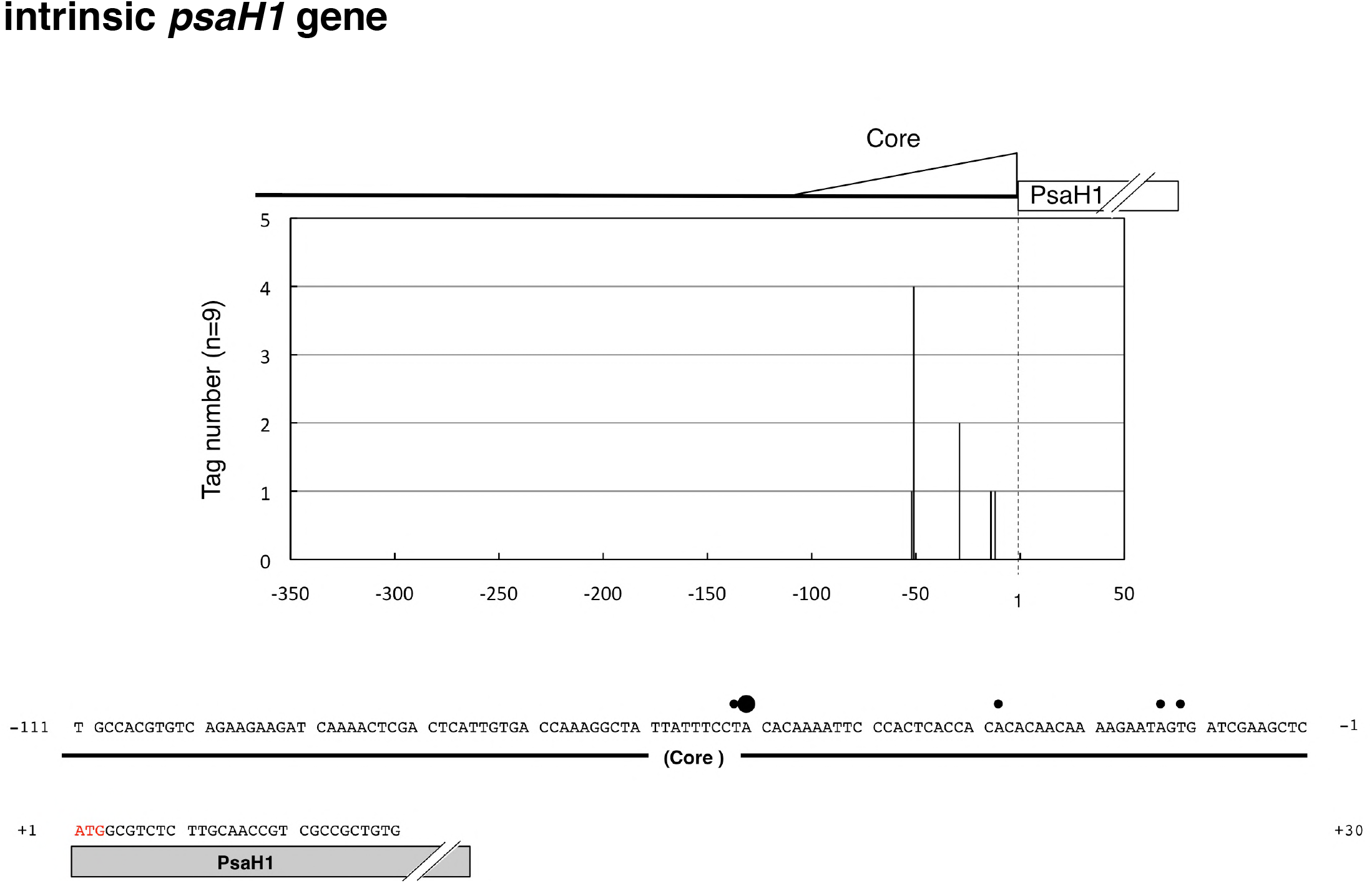
TSS distribution profile of *psaH1* of *Arabidopsis thaliana*. ATG in red indicates the translation start site (+1) of *psaH1*.psaH1::LUC

**Figure S4.**
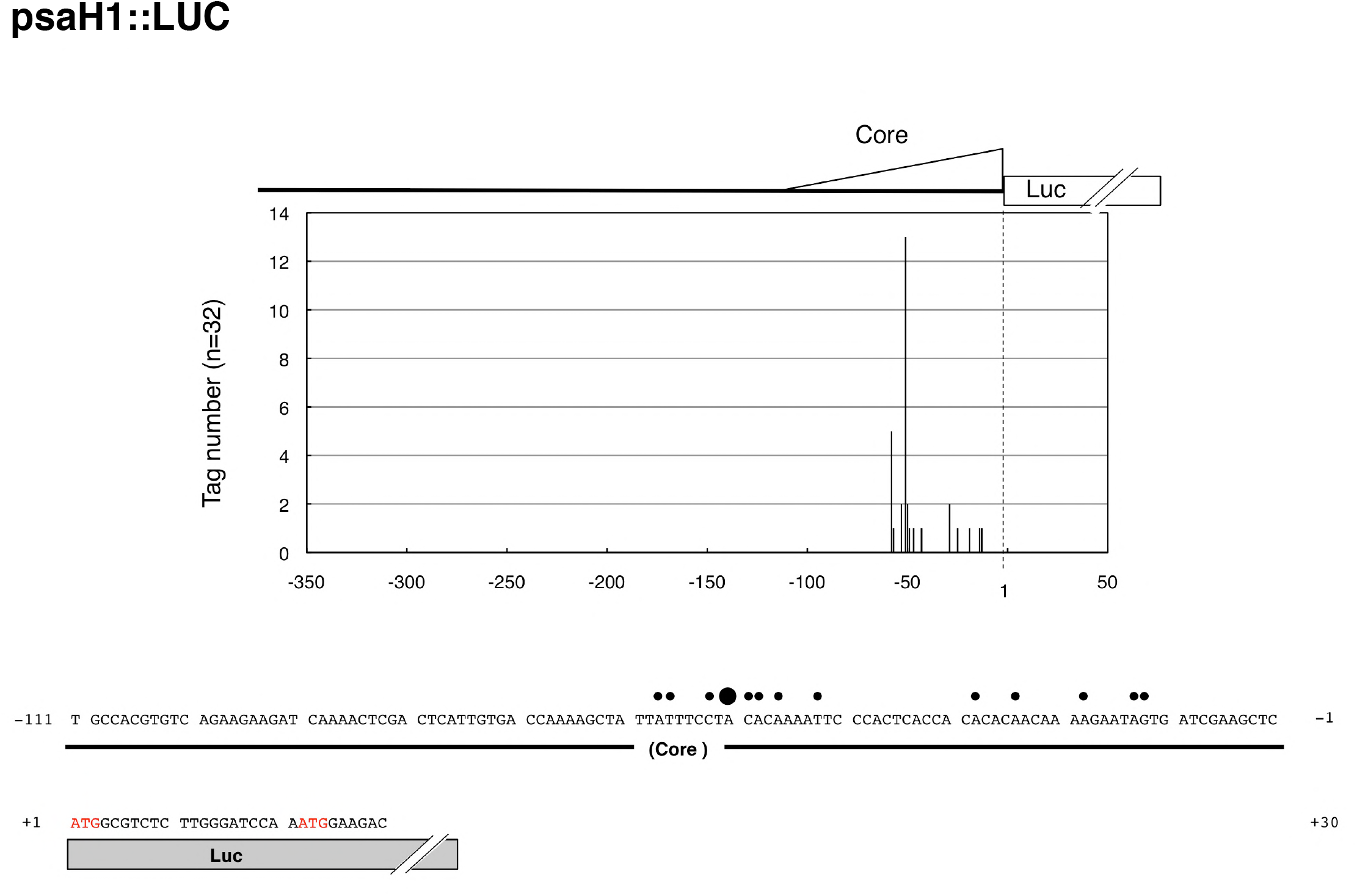
TSS distribution profile of PsaH1::LUC. Whole promoter region plus translation initiation context of *psaH1* (−2000 to +12 relative to the ATG initiation codon) was translationally fused with the firefly luciferase (LUC) gene via a *BamHI* linker. ATG initiation codons (red) of *psaH1* and LUC are in frame, but are separated by 18 bp.

**Figure S5.**
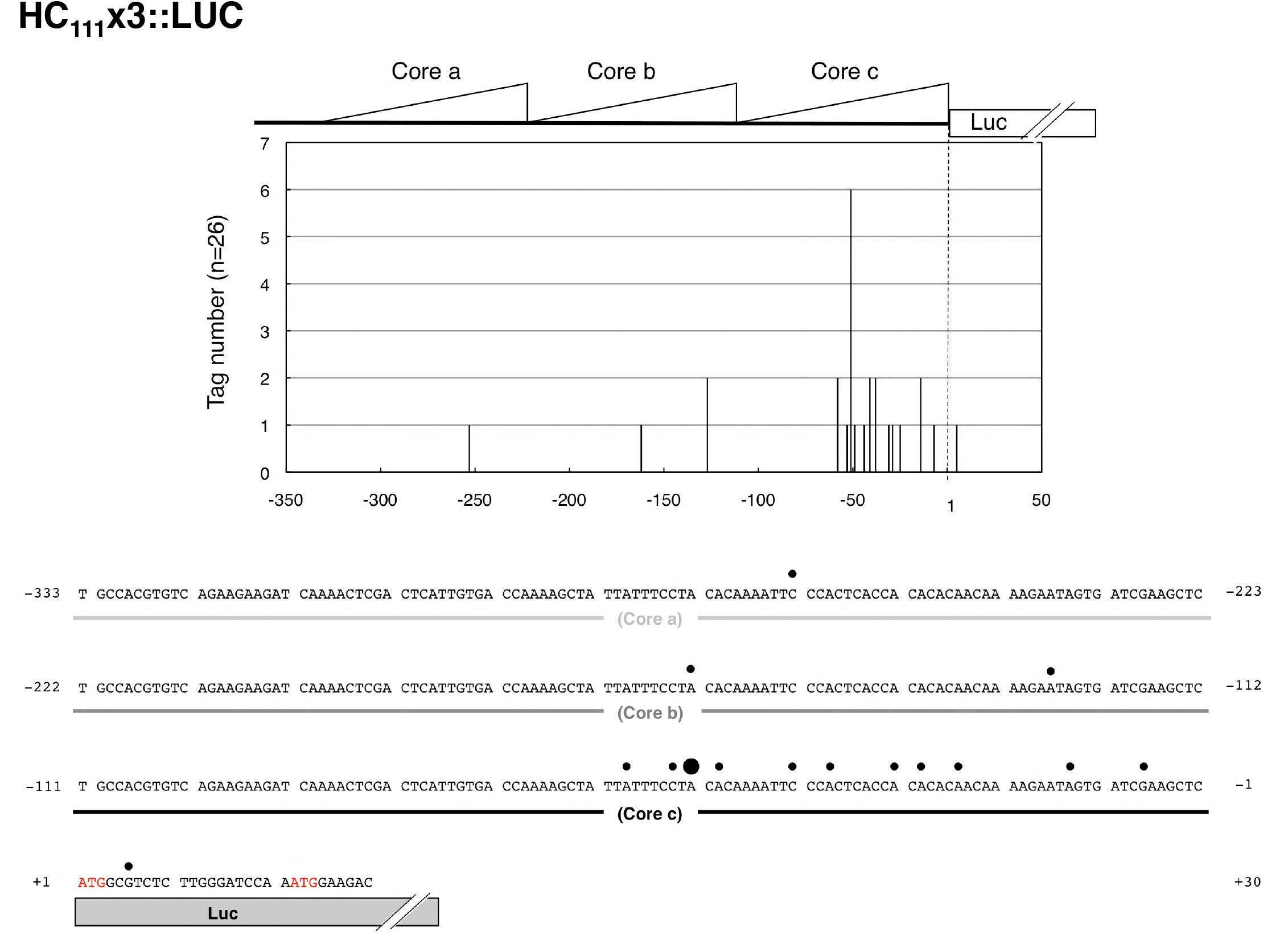
TSS distribution profile of HC_111_x3::LUC. TSSs occur preferentially in segment c.

**Table S1.**
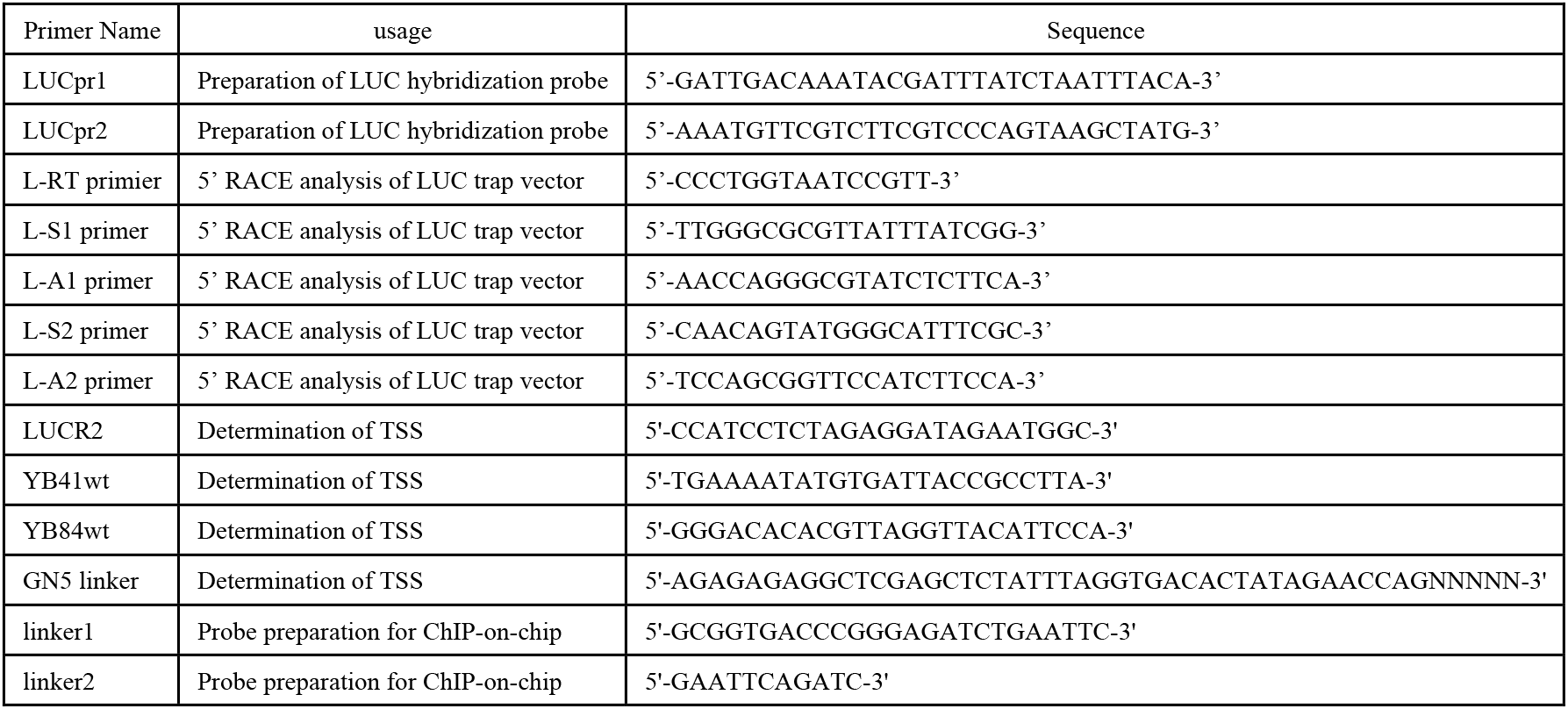
Miscellaneous primers.

**Table S2.**
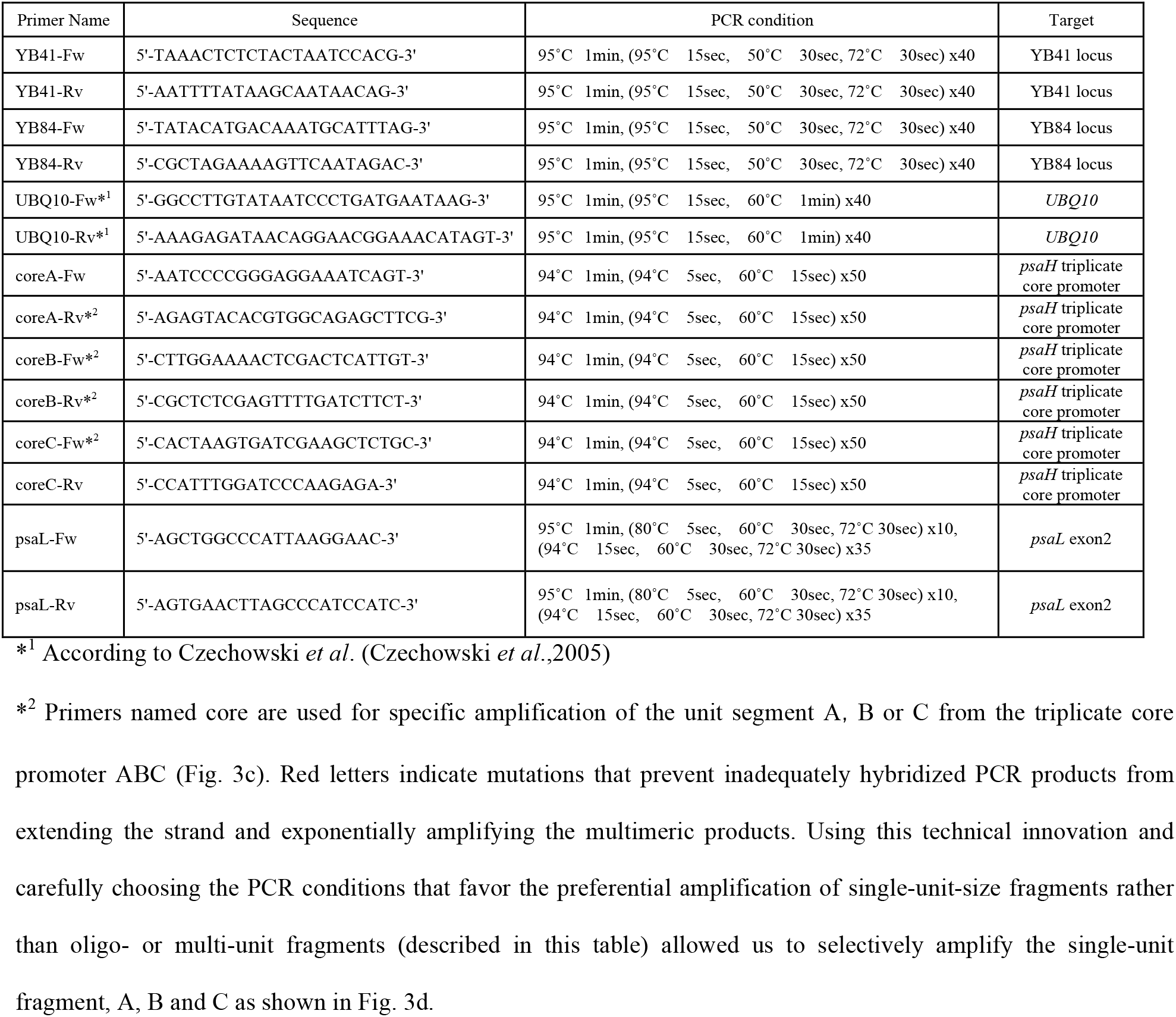
Primers and thermal cycling conditions of real-time PCR analysis.

